# SPATIO-TEMPORAL DYNAMICS OF EARLY SOMITE SEGMENTATION IN THE CHICKEN EMBRYO

**DOI:** 10.1101/2023.04.05.535703

**Authors:** Ana Cristina Maia-Fernandes, Ana Martins-Jesus, Tomás Pais-de-Azevedo, Ramiro Magno, Isabel Duarte, Raquel P. Andrade

**Affiliations:** ABC-RI, Algarve Biomedical Center Research Institute, Faro, Portugal; Faculdade de Medicina e Ciências Biomédicas (FMCB), Universidade do Algarve, Campus de Gambelas, 8005-139 Faro, Portugal; Pattern Institute, Faro, Portugal; CINTESIS@RISE, Universidade do Algarve, 8005-139 Faro, Portugal; Champalimaud Research Program, Champalimaud Center for the Unknown, Lisbon, Portugal

**Keywords:** somitogenesis, *hairy1*, embryo clock, morphometrics, periodicity

## Abstract

During vertebrate embryo development, the body is progressively segmented along the anterior-posterior (A-P) axis early in development. The rate of somite formation is controlled by the somitogenesis embryo clock (EC), which was first described as gene expression oscillations of *hairy1* (*hes4*) in the presomitic mesoderm of chick embryos with 15-20 somites. Here, the EC displays the same periodicity as somite formation, 90 min, whereas the posterior-most somites (44-52) only arise every 150 minutes, matched by a corresponding slower pace of the EC. Evidence suggests that the rostral-most somites are formed faster, however, their periodicity and the EC expression dynamics in these early stages are unknown. In this study, we used time-lapse imaging of chicken embryos from primitive streak to somitogenesis stages with high temporal resolution (3-minute intervals). We measured the length between the anterior-most and the last formed somitic clefts in each captured frame and developed a simple algorithm to automatically infer both the length and time of formation of each somite. We found that the occipital somites (up to somite 5) form at an average rate of 75 minutes, while somites 6 onwards are formed approximately every 90 minutes. We also assessed the expression dynamics of *hairy1* using half-embryo explants cultured for different periods of time. This showed that EC *hairy1* expression is highly dynamic prior to somitogenesis and assumes a clear oscillatory behaviour as the first somites are formed. Importantly, using *ex ovo* culture and live-imaging techniques, we showed that the *hairy1* expression pattern recapitulates with the formation of each new pair of somites, indicating that somite segmentation is coupled with EC oscillations since the onset of somitogenesis.

**Highlights:**

- Time of early somite formation can be inferred from sequential length measurements
- The cranial-most somites are formed faster than trunk somites
- Oscillations of *hairy1* expression are temporally coupled with early somite formation

**GRAPHICAL ABSTRACT:** 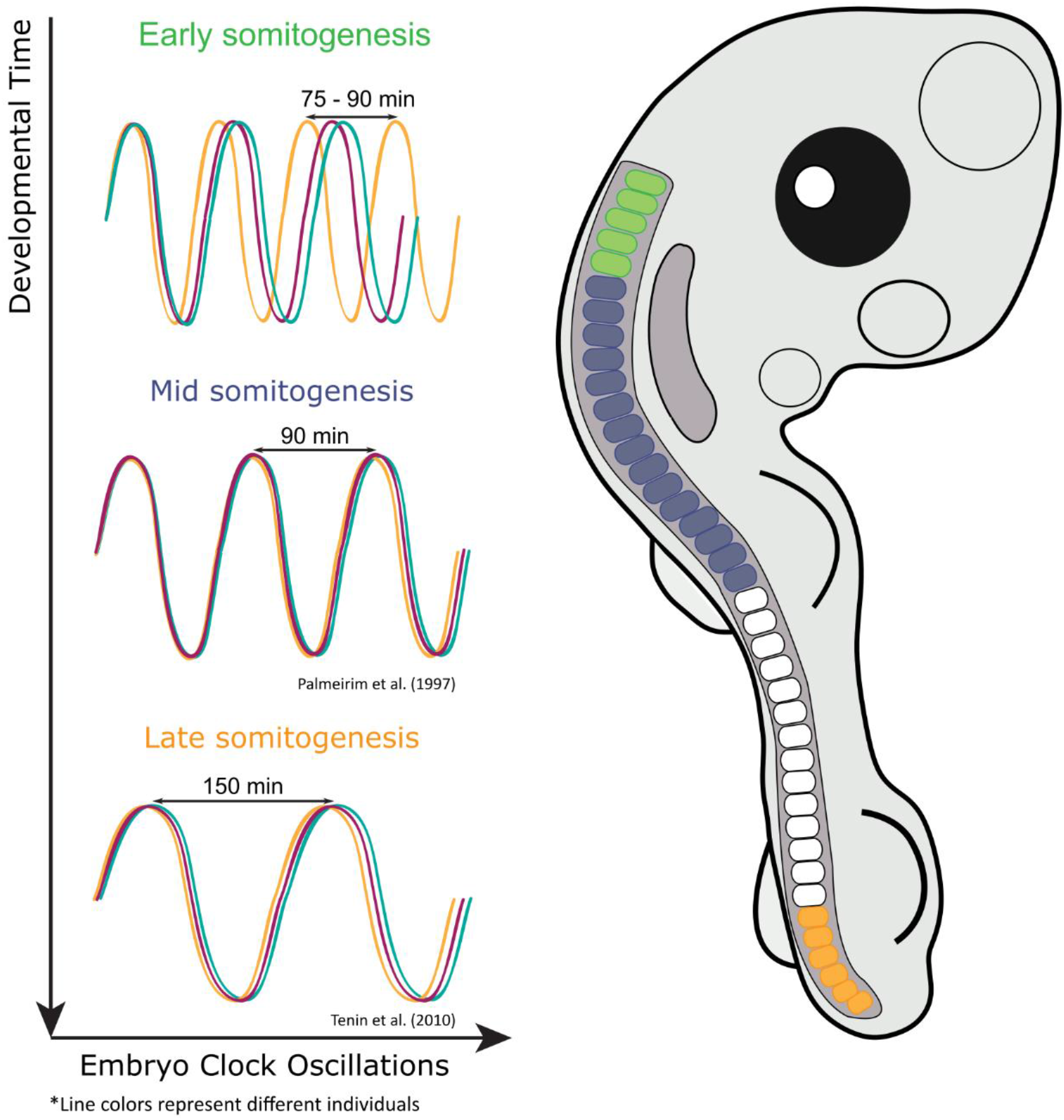

## INTRODUCTION

During embryo development the molecular and cellular events leading to the formation of a functional organism need to occur in the correct order and at the right pace. Vertebrate embryo body segmentation along the antero-posterior (A-P) axis is a paradigmatic example of an ordered, time-controlled process. During somitogenesis, the anterior-most presomitic mesoderm (PSM) is periodically segmented into somites which are the precursor metameric structures of the axial skeleton and musculature. Somites are formed sequentially along the embryo A-P body axis where they will give rise to different adult structures (Bodo Christ & Ordahl, 1995). In the chick embryo, the first four-five somites will originate occipital structures, somites 5-19 give rise to cervical vertebrae, somites 19-26 have thoracic identity, and somites 26-30, 30-39, and 39-52 have lumbar, sacral, and coccygeal identities, respectively (Christ et al., 2000; Christ & Ordahl, 1995; Huang et al., 2000).

The periodicity of morphological somite formation is controlled by the somitogenesis embryo clock (EC), comprising genes with cell-autonomous oscillatory expression in the PSM driven by negative feedback loops (reviewed in Carraco et al., 2022). Chicken *hairy1* (*hes4*), a member of the Hairy and Enhancer of split (HES) protein family of transcription factors, was the first EC gene described, oscillating in the PSM every 90 minutes, the same periodicity as somite formation (Palmeirim et al., 1997). The process of somitogenesis has been studied focusing mainly on the late-cervical/thoracic somites, where the temporal dynamics of somite formation and underlying EC oscillations are well known. However, previous work showed that the tempo of somite formation varies along the embryo A-P axis (Schröter et al., 2008; Schubert et al., 2001; Tam, 1981; Tenin et al., 2010). In the chicken embryo, the posterior-most somites (44-52), are formed at a slower pace – a new pair arises only every 150 minutes. Importantly, this is matched by a corresponding slower pace of EC oscillations (Tenin et al., 2010). The time of somite formation in the zebrafish tail has also been reported to increase over development (Kimmel et al., 1995; Schröter et al., 2008). In fact, the cranial-most somites form at a faster rate, as also seen in mouse and amphioxus (Schubert et al., 2001; Tam, 1981). In the chicken embryo, the first somites were described to arise almost simultaneously (Dias et al., 2014), raising the hypothesis that the EC may have very different expression dynamics during occipital somite formation.

The somitogenesis clock genes *lunatic fringe (lfng), hairy1 (hes4)*, and *hairy2 (hes1)* are dynamically expressed since gastrulation in the chicken embryo (Jouve et al., 2002) and early somitogenesis chicken embryos display different patterns of EC gene expression in the PSM, similarly to what is observed in later stages (Rodrigues et al., 2006). However, the temporal dynamics of EC gene expression during early somitogenesis remains unknown, and it is unclear whether the formation of the early-most somites is coupled to the operation of the segmentation clock. In this work, we used time-lapse imaging of the chick embryo to make a thorough characterization of the spatial and temporal properties of the formation of the cranial-most somites. By assessing *hairy1* gene expression in different time points of early developmental stages, we evidence that EC periodicity is coupled with somite formation rate since the onset of somitogenesis.

## METHODS

### Embryos and culture conditions

Fertilized chicken eggs were obtained from commercial sources (Pintobar Exploração Agrícola, Lda, Portugal) and incubated at 38ºC in a humidified atmosphere up to the desired developmental stage (Hamburger & Hamilton, 1992). For *in vivo* imaging HH4 to HH10 chicken embryos were cultured using the Easy Culture system (Chapman et al., 2001) with the ventral side facing up, in a humidified stage-top incubator (UNO-T H-301-k, OKOLAB) at 38ºC.

### Image acquisition and analysis

Time-lapse movies performed with 3-min interval resolution, and in situ hybridization photos were acquired using a Zeiss SteREOLumarV12 stereomicroscope coupled with a Zeiss Axiocam Mrc camera. Images were adjusted for brightness and contrast using the free software Gimp (v. 2.10.8).

### Somite nomenclature

Somites were numbered according to their craniocaudal position along the embryo’s body axis, so that the somite number always corresponds to the same morphological segment, irrespective of the developmental stage of the embryo analysed.

### Embryo measurements

Length measurements were performed in all acquired frames of somitogenesis staged embryos, using Axiovision Se64 Rel 4.9.1 (Carl Zeiss) or Zen 2.5 (blue edition, Carl Zeiss). Measurements were taken from the anterior border of the first somite until the last completely formed cleft. A somitic cleft was considered completely formed only when it was clearly visible from its axial to lateral limits. We conducted a total of 2836 measurements on 13 different chicken embryos in stages HH7 to HH10 (corresponding to somites 1 to 10) and HH11+ to HH13+ (corresponding to somites 14 to 20). To minimize measurement biases, at least two different operators performed independent measurements.

### Data analysis

Data analysis was performed using R (version 3.5.2) (R Core Team, 2017) in RStudio (Version 1.1.463) (R Studio Team, 2015). R packages ggplot2 (version 3.1.1) and dplyr (version 0.8.0.1) were used for data tidying, wrangling, and visualization. The raw data of the measurements performed is deposited in Figshare (10.6084/m9.figshare.22341070), and the analysis algorithm has been encapsulated into the open-source R Package SomiteExplorerR, freely available in GitHub (github.com/iduarte/SomiteExplorerR).

To determine the period of early somite formation, we analysed consecutive measurements (every 3 minutes) of the total length of the embryo segmented region (SEG). Data analysis entailed three major steps: (i) the automated classification of somites by length increment, (ii) the calculation of each somite length, and (iii) the somite period inference. Our algorithm automatically assigns the somite number based on consecutive differences between total lengths. Briefly, a length difference between two consecutive measurements greater than 50 µM (value empirically derived) was used to signal the formation of a new somite. This allows the automated classification of the somite number pertaining to each of the 2836 measurements taken, without the need to add these data manually (which is time consuming and error-prone). Simultaneously, the calculation of the differences between total segmented length enables the estimation of the average length of each somite. The first measurement of SEG corresponds to the length of the first somite, so this approach allows the inference of the time of formation from the second somite onwards. The calculation of the period of formation of somite n was performed by subtracting the first time point of somite n to the first time point of somite n-1. Further details can be found in the SomiteExplorerR package repository.

### Embryo explant culture

Embryo explant cultures were performed as previously described (Palmeirim et al., 1997). Briefly, the embryo caudal portion was bisected along the midline including neural tube and notochord. Each isolated explant was placed on an Isopore Membrane Filter 0,8μm (Millipore) floating on M199 medium supplemented with 10% chick serum, 5% fetal bovine serum and 100 U/ml of penicillin/streptomycin and incubated at 37ºC and 5% CO_2_. After 5-10 min, one of the explant halves was immediately fixed, while the other was cultivated further for different periods of time.

### *In situ* hybridization

*In situ* hybridization was performed as previously described for *hairy1* (Palmeirim et al., 1997) and *hairy2* (Jouve et al., 2000).

## RESULTS

### Time of somite formation can be directly inferred from length increments in the embryo segmented region

The time required for the formation of each of the first somites in the chicken embryo is unknown. To thoroughly assess the dynamics of this process, we performed time-lapse imaging of developing chick embryos since primitive streak stages (HH4-6) until 10-somite stage (HH10), with 3-minute intervals (Figure 1A). We carefully examined all acquired frames for each embryo and measured the A-P distance between the first and last completely formed somitic clefts, hereafter referred to as the segmented region (SEG) (Figure 1B). An automated algorithm was developed to calculate the length differences between measurements, evidencing that SEG length remained approximately constant in multiple consecutive frames, intercalated with periodic bursts of length increments, corresponding to the addition of each new somite at the posterior end of SEG (Figure 1C,D). Formation of each new somite was automatically detected by the algorithm when the SEG increment exceeded an empirically derived threshold (in this case, 50μm; Figure 1C), and a somite-specific identity was assigned (Figure 1D). Note that the measurements included only the whole segmented region of the embryo, meaning that the raw data did not contain any information regarding the number of somites therein. We applied this methodology to a total of 11 chick embryos in early somitogenesis stages and four embryos encompassing the formation of somites 15-20 (Supplementary Figure 1), where the somitogenesis properties are best described. The results are summarized in Table 1 and Supplementary Table 1. This approach provided direct information on the length (L, Figure 1D; Table 1) of each newly formed somite, corresponding to the length increment observed at each peak (Figure 1C). Moreover, it allowed us to infer *time* from *space*, since the time elapsed between two consecutive SEG length increments corresponds to the time required to form each new somite (T, Figure 1D; Table 1).

**Figure 1.**
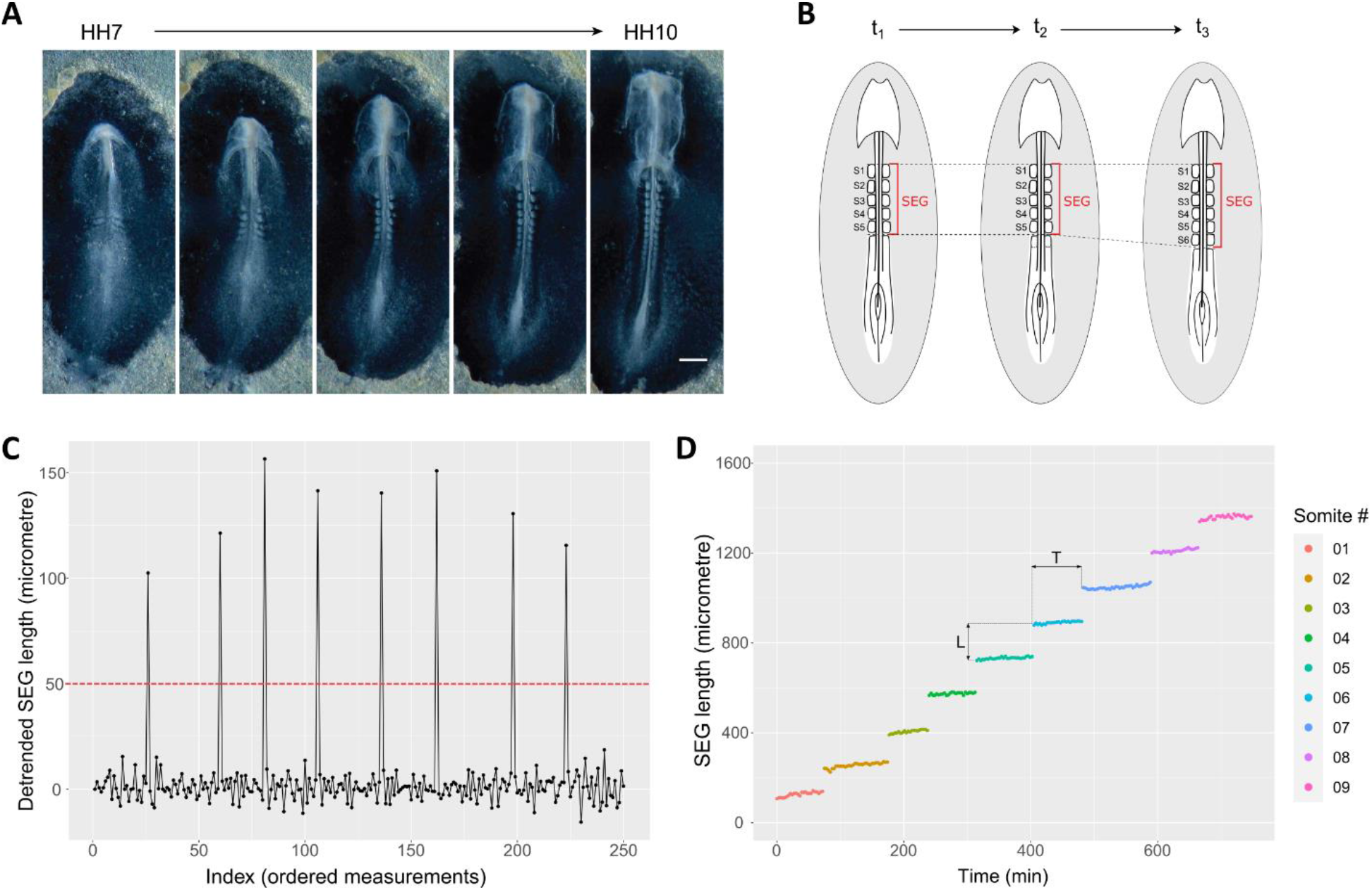
Methodological approach to determine somite length and time of formation in the chick embryo. **(A)** Representative images of time-lapse visualization of early chick embryo development, encompassing the formation of the first ten somites. Scale bar: 0.5 mm; **(B)** Graphical representation of the measurements performed to determine the length of the embryo segmented region (SEG); **(C)** The formation of each new somite was automatically detected as peaks of SEG length increments, using a custom-made data detrending algorithm with 50 μm as cut-off. Data from a single embryo is presented; **(D)** SEG length of early chicken embryos over time. Data from a single embryo is presented. L: somite length; T: time of somite formation.

**Table 1.** Length and Time of formation of chick early somites. Mean ± standard deviation is shown.

| Somite # | Embryos sampled (n) | Length ($\mu\text{m}$) | Time of Formation (min) |
| --- | --- | --- | --- |
| 1 | 11 | $117.57 \pm 15.97$ | NA |
| 2 | 11 | $127.39 \pm 8.84$ | $60.55 \pm 23.19$ |
| 3 | 11 | $146.65 \pm 18.59$ | $78.27 \pm 19.51$ |
| 4 | 11 | $154.5 \pm 17.72$ | $72.55 \pm 20.43$ |
| 5 | 11 | $147.78 \pm 19.29$ | $76.91 \pm 12.61$ |
| 6 | 11 | $139.57 \pm 18.79$ | $87.27 \pm 13.78$ |
| 7 | 9 | $138.01 \pm 22.07$ | $80.67 \pm 23.5$ |
| 8 | 6 | $145.74 \pm 22.07$ | $91.5 \pm 14.79$ |
| 9 | 5 | $140.22 \pm 27.72$ | $85.2 \pm 11.92$ |
| 14 | 4 | $115.92 \pm 16.83$ | NA |
| 15 | 4 | $130.36 \pm 12.63$ | $88.5 \pm 13.53$ |
| 16 | 4 | $142.53 \pm 11.68$ | $81.75 \pm 13.94$ |
| 17 | 4 | $171.14 \pm 23.8$ | $89.25 \pm 7.89$ |
| 18 | 4 | $191.14 \pm 32.72$ | $96 \pm 14.7$ |
| 19 | 4 | $177.61 \pm 58.62$ | $85.5 \pm 14.39$ |
| 20 | 3 | $156.22 \pm 21.09$ | $81 \pm 6$ |

NA: not applicable; corresponds to the starting point of the time-lapses, for which the somite formation time cannot be calculated using our method.

### Anterior-posterior length of rostral somites

A direct output of our analysis is the length of each individual somite at the moment of formation (Table 1, Figure 2, Supplementary Figure 2). We found that somite average length ranges from ∼118 - 191 μm (somites 1 and 18 respectively). Somite length of the first nine somites did not vary considerably, confirming that the appearance of each new cleft corresponded to the formation of a single somite. The large variability in measurements of somites 17-20 most probably results from the rotation of the embryo body in these developmental stages.

**Figure 2.**
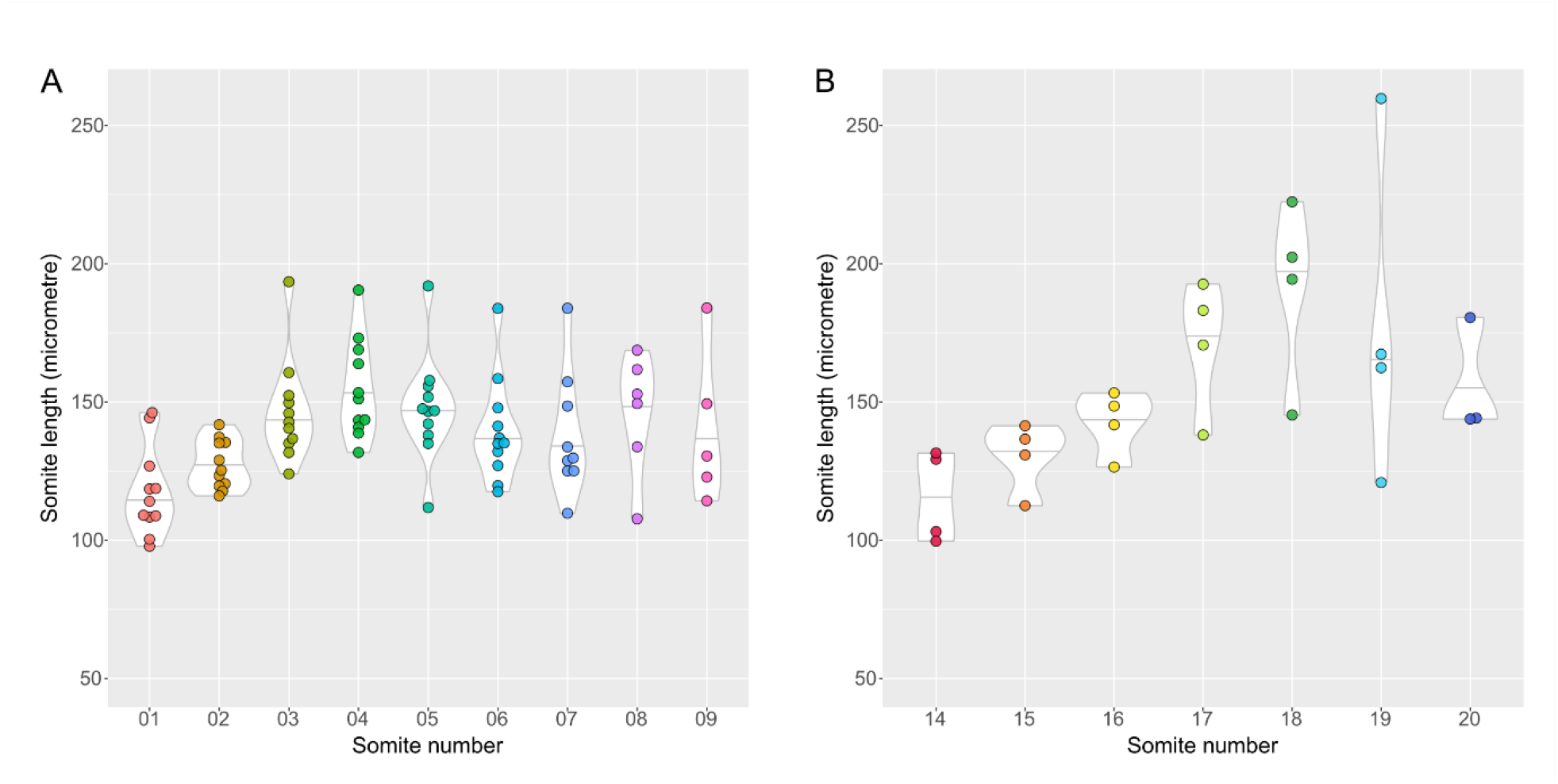
Somite length upon segmentation at different axial positions. **(A,B)** Violin-plot distribution of the observed lengths of individual somites 1-9 (A) and 14-20 (B).

### Occipital somites are formed faster than cervical and trunk somites

A clear observation from our data is that all somites are formed consecutively, i.e. one after the other (Figure 3A, B and Supplementary Figure 1 A, B). In no case did we observe the simultaneous formation of more than one of the early-most somites (n=11; Supplementary Figure 1A), as was previously suggested (Dias et al., 2014). We did find, however, that the first five somites, corresponding to the occipital precursors, form faster, at an average rate of 75.57±3.21 min/somite (Figure 3A). Somites 6-9 then form at a rate of 91.95 ± 12.24 min/somite (Figure 3A). This closely matches the rate of formation of somites 15-20, of 87.02 ± 3.99 min/somite (Figure 3B, Table 1), which is approximately 90 minutes, as previously described using other experimental strategies (Palmeirim et al., 1997).

**Figure 3.**
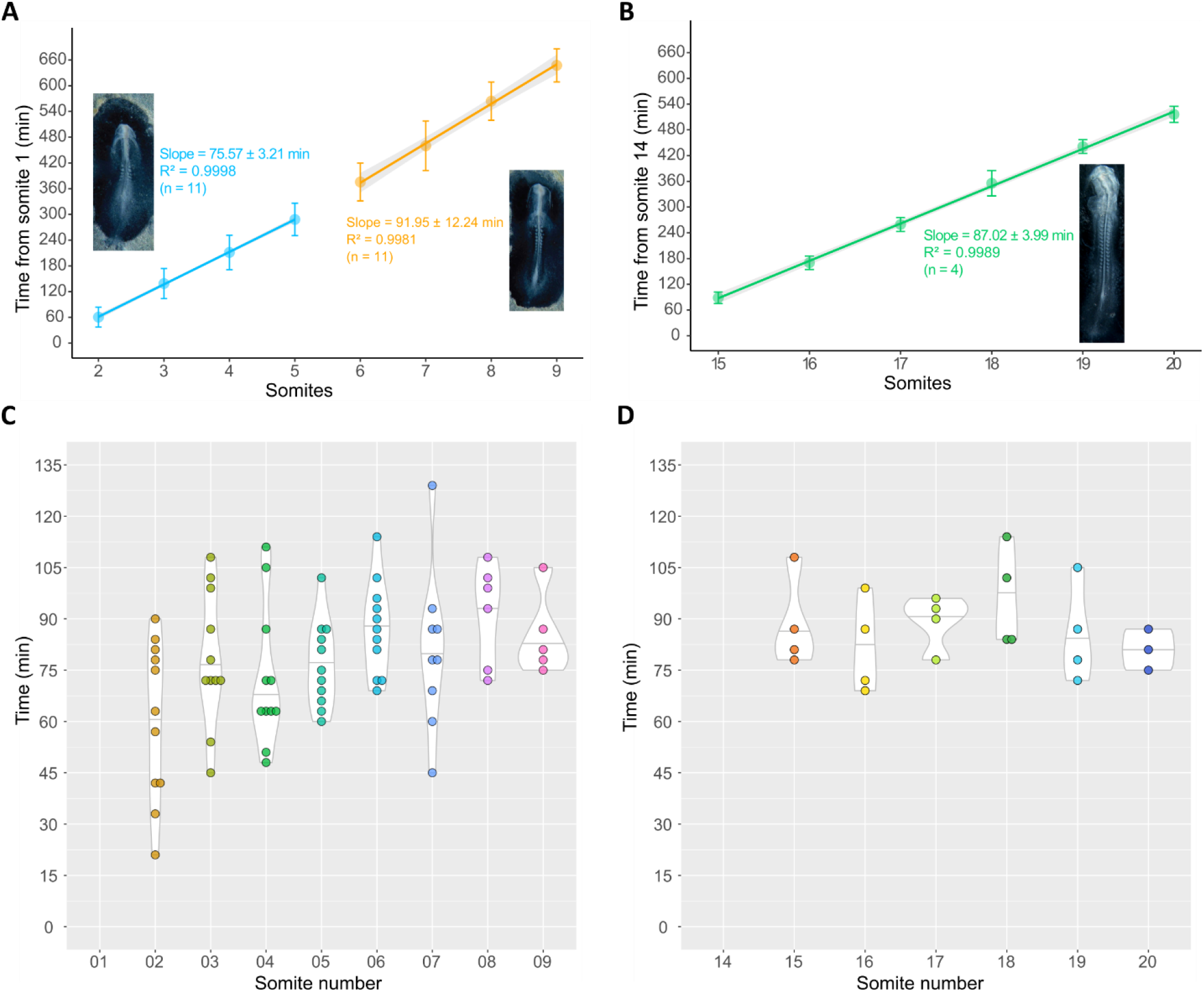
Somitogenesis period at different axial positions inferred from length increments of the embryo segmented region. **(A,B)** Mean somite formation time ± SD of individual somites 2-9 (A) and 15-20 (B). The somitogenesis period of each group of somites was estimated as the slope of the linear regression with 95% confidence interval. A linear increase in somite number over time is evident. The occipital somites are formed faster than the neck and trunk somites. Insets are representative images of embryos in the developmental stages analysed, positioned ventral side up. **(C**,**D)** Violin-plot distribution of the observed periods of individual somites 2-9 (C) and 15-20 (D). The early-most somites have higher variability of somite formation time. The period for somites 1 and 14 is not displayed since it cannot be inferred with our methodology. SD: standard deviation.

Remarkably, there is substantial variability in the time of formation of the early-most somites (Figure 3C, Supplementary Figure 3), which gradually stabilizes until somite 8 onwards, where both somite formation time and variability is equivalent to that observed for somites 15-20 (Figure 3C, D; Table 1). Altogether, we report that the occipital somites form faster and with greater temporal variability than the neck and trunk somites.

### *hairy1* gene expression oscillates concomitantly with the formation of the first somites

Morphological somite formation is described to be preceded by oscillations of EC gene expression in the PSM with the same periodicity (Palmeirim et al., 1997; Tenin et al., 2010). To evaluate if this holds true for the occipital somites, we characterized the expression patterns of *hairy1* and *hairy2* before the onset of somitogenesis (Figure 4A, B), and in early somitogenesis stages (Figure 5A,B) using *in situ* hybridization. We observed that both *hairy1* and *hairy2* present different expression patterns in embryos within the same developmental stage, which is a distinctive feature of EC operation (Palmeirim et al., 1997). In primitive streak-embryos, *hairy1* expression presented highly variable patterns along the A-P axis of the primitive streak and was always observed in the Hensen’s node (n=57) (Figure 4A). *hairy2* expression was also very dynamic along the embryo A-P axis (n=20) (Figure 4B), as previously described (Jouve et al., 2002).

**Figure 4.**
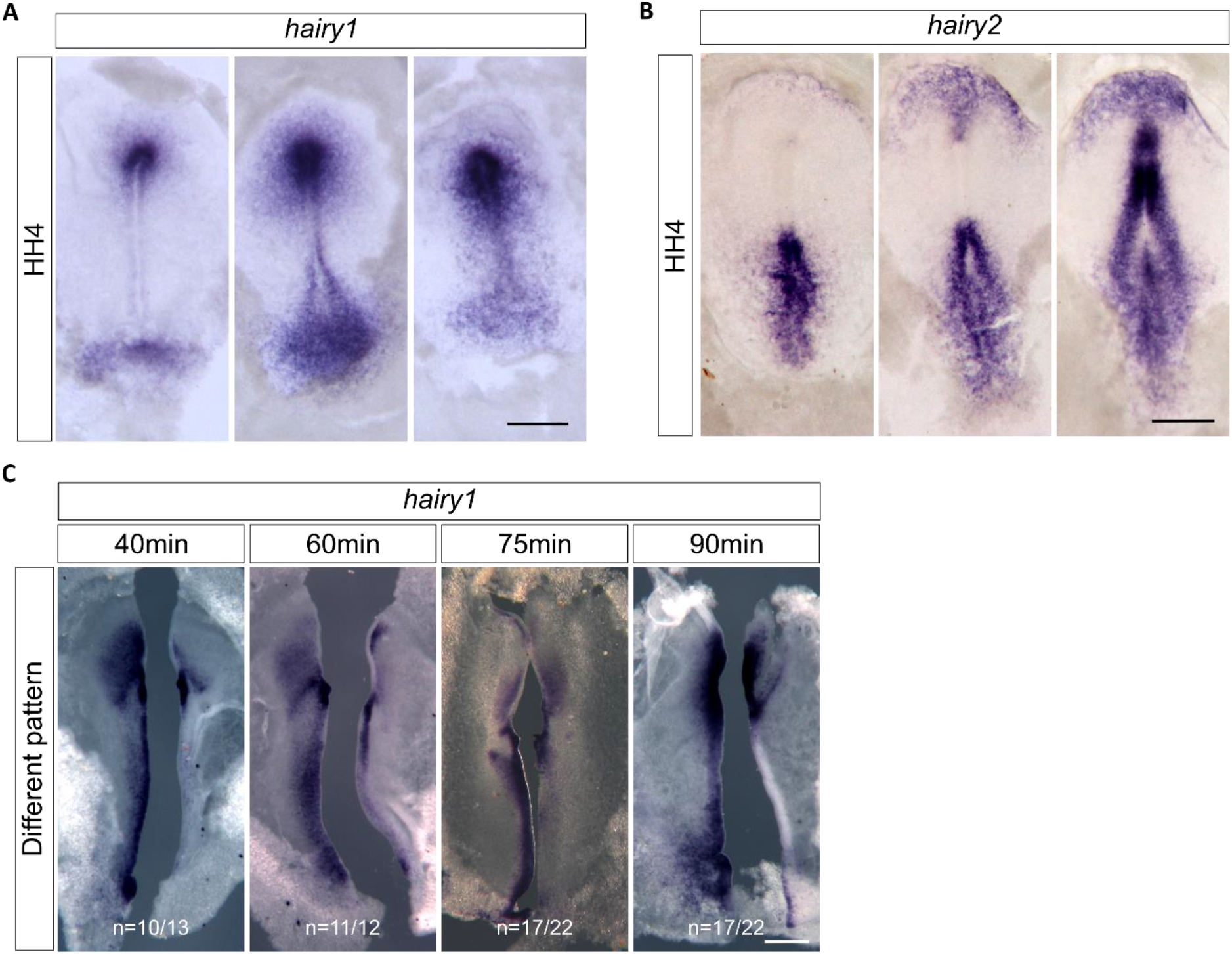
Expression dynamics of *hairy1* and *hairy2* in the early chick embryo. *In toto in situ* hybridization for *hairy1* **(A)** and *hairy2* **(B)**. Embryos within the same morphological stage present very different expression patterns; **(C)** Different *hairy1* expression pattern observed in HH4-HH6 embryo halves cultured for different periods of time. Scale bars: 0.5 mm.

**Figure 5.**
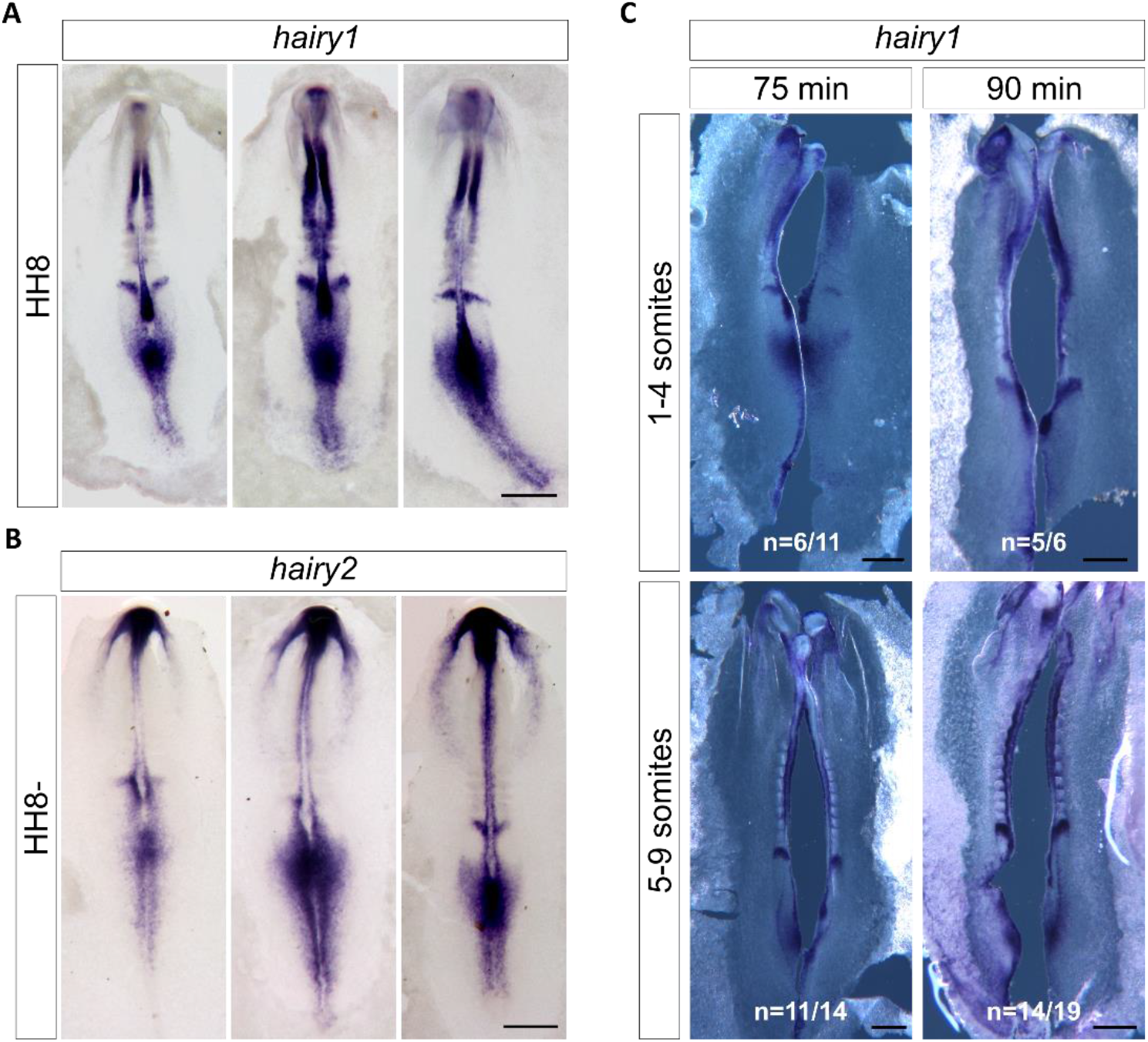
Expression dynamics of *hairy1* and *hairy2* in early somitogenesis stages. *In toto in situ* hybridization for *hairy1* **(A)** and *hairy2* **(B)**. Embryos within the same morphological stage present different expression patterns; **(C)** *hairy1* expression pattern is recapitulated in embryos halves cultured for 75 - 90 minutes. Scale bars: 0.5 mm.

To determine if *hairy1* expression is oscillatory, i.e. if the expression pattern is recapitulated over time, we cut the embryo along the midline, fixed one half and cultured the other half for additional time periods. Then, *in situ* hybridization was performed simultaneously in both explants and the expression patterns obtained were compared. When this procedure was applied to early gastrulating embryos (HH3+ to HH6), different *hairy1* expression patterns were obtained in all the incubation times tested (Figure 4C). A 40-minute incubation was enough to drastically change *hairy1* expression and, although up to 20% of the explants presented the same expression pattern, a consistent recapitulation of *hairy1* expression was not obtained in any of the time points tested (Figure 4C). As somitogenesis takes place, *hairy1* and *hairy2* expression patterns retain their dynamic properties (Figure 5A, B) (Rodrigues et al., 2006). But, in this case, recapitulation of *hairy1* expression patterns was observed after 75-90 minutes of incubation (Figure 5C), as soon as the first somite was formed.

Together, our data evidences that *hairy1* expression is very dynamic since early gastrulation, and that clear oscillations of EC gene expression are established as the first somites are formed.

### Periodicity of *hairy1* oscillations is coupled to the rate of early somite segmentation

The rate of trunk somite formation corresponds to the periodicity of EC cyclic expression in the PSM (Palmeirim et al., 1997) and, as the periodicity of somite formation is decelerated towards the caudal embryo, it consistently matches the periodicity of EC gene expression oscillations (Tenin et al., 2010). Here, we asked if the EC dynamics could also be coupled to the formation of the first somites. Since the time of formation of the occipital somites is highly variable (Figure 3C), we employed an experimental approach that allowed us to assess the expression pattern of *hairy1* at the precise moment of the formation of two consecutive somitic clefts in a single embryo (Figure 6). Our strategy consisted in performing live-imaging of the early embryo until a new somitic cleft was formed. At this moment (t0), we surgically removed and immediately fixed the PSM on one side of the embryo, encompassing the last formed somite until the Hensen’s node. Live imaging was quickly resumed until the formation of the next somitic cleft in the contralateral side (tf), and the remaining embryonic tissue was fixed. Then, both tissues were jointly processed for *in situ* hybridization (Figure 6). We found that the expression pattern that *hairy1* presented when the initial somite cleft appeared (t0) was recapitulated at the moment of formation of the next consecutive cleft (tf). This was true both for the formation of occipital somites (somites 2-4, n=3/4) and early cervical somites (somites 6-7, n=2/3) (Figure 6). These results show that *hairy1* oscillations and somite formation dynamics are already coupled in the rostral-most somites, suggesting that the EC may underlie the pace of somite formation along the entire embryo A-P axis.

**Figure 6.**
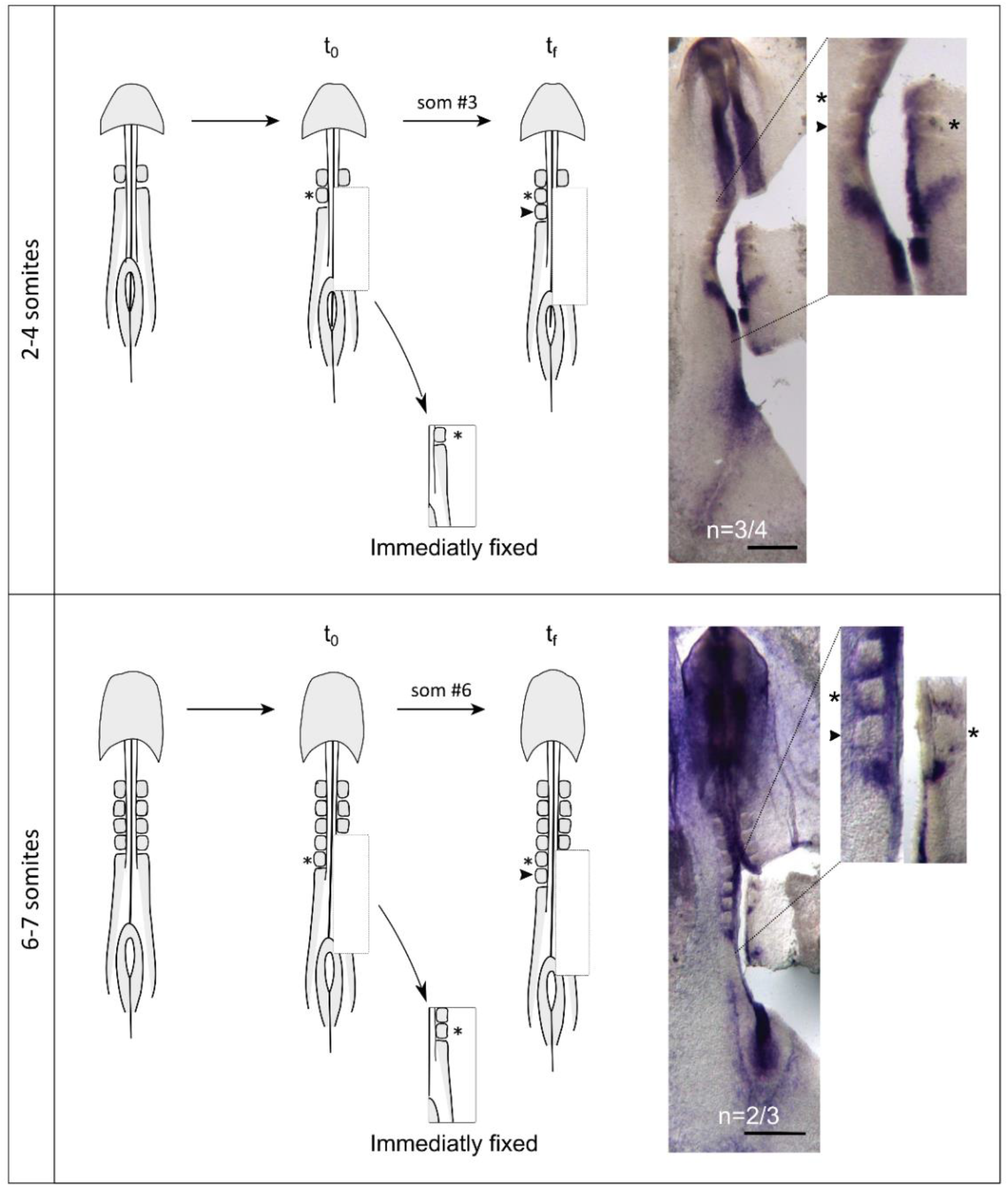
*hairy1* expression oscillates with the same periodicity as early somite formation. Live-imaging of early embryos was performed to identify the moment of formation of a new somitic cleft. The PSM tissue was excised from one side and immediately fixed (t0), while the remaining tissue was re-incubated until the formation of the next somitic cleft (tf). The same *hairy1* expression pattern was obtained by *in situ* hybridization in both halves of embryos with 2-4 somites (n=3/4) and 6-7 somites (n=2/3), evidencing that cyclic expression of *hairy1* is temporally coupled to the formation of a new somite. Asterisks and arrowheads mark the last formed somite at t0 and at tf, respectively. Scale: 0.5 mm.

## DISCUSSION

The spatio-temporal characteristics of somite formation vary considerably along the vertebrate embryo A-P axis. Although it is well known that the periodicity of EC oscillations underlies the rate of formation of the late cervical, trunk, and caudal somites, this has not been established for the rostral-most somites. Here, we present a thorough characterization of the size and time of formation of the first ten somites in the chicken embryo. We employed a novel strategy to infer somite formation time from length measurements, and report that the early-most somites form at a faster rate than their later counterparts. We also observed that the somitogenesis EC is temporally coupled with somite formation since the onset of embryo body segmentation.

### Inferring *time* from *space*: characterization of early somite formation

The rostral-most somites have previously been described to form very rapidly in the chicken embryo (Dias et al., 2014). Accordingly, this process must be analysed with high temporal and spatial resolution to properly assess the dynamics of the beginning of somitogenesis. Live-imaging enables the study of morphological events in the same embryo throughout a large temporal window, allowing the distinction between the variability of a particular process from the variability inherent to a population of different samples.

Our strategy to characterize the dynamics of early chick somite formation relied on time lapse imaging of single embryos with high temporal sampling (3-minute intervals). Then, using a single measured variable – the length of the embryo’s total segmented region – we computationally inferred the somite identity, length, and formation time using the calculated length increments. This approach presents important advantages, namely, (1) the error originated from sample variability is minimized by analysing the formation of successive somites in the same embryo over time, allowing to derive conclusions more confidently using a smaller number of biological samples; (2) somite size and formation time are derived simultaneously from a single measurement, thus avoiding two distinct errors associated with the measuring instruments (as when separately using a ruler and a clock); and, (3) the algorithm developed for automated data analysis assures reproducibility while allowing a significant increase in the number of samples analysed, therefore improving the precision of future large-scale studies.

### Spatio-temporal properties of the rostral somite segmentation

Application of our methodology unveiled the initial length and time of formation of the rostral-most somites of the chicken embryo. The early somite average length ranged from approximately 118 µm to 155 µm, which is in line with previous work (Herrmann et al., 1951).

The first five somitic clefts were formed sequentially over time, with an average periodicity of 75 minutes. This detailed live-imaging approach, thus, clarifies that the first somites in the chicken embryo are not formed simultaneously, as previously suggested (Dias et al., 2014).

Our work complements previous studies that reported an increase in somite formation time along the embryonic A-P axis (Schröter et al., 2008; Schubert et al., 2001; Tam, 1981; Tenin et al., 2010). We show that the first five somites form with an average periodicity of 75 min, after which somitogenesis gradually stabilizes at the previously reported 90-minute rate (Palmeirim et al., 1997). Finally, the last somites form every 150 minutes (Tenin et al., 2010).

### The Embryo Clock in early somitogenesis

The fact that we observed sequential formation of the first somites, and that somite clefts are regularly spaced originating somites of similar sizes, strongly suggests that a “clock-and-wavefront” type mechanism could already be operating in these early stages. Previous studies suggested that the EC is dispensable for early somite formation (Dias et al., 2014). In fact, inhibition of BMP signalling in the chick epiblast is sufficient to allow rosette-shaped somites to form, without perceivable dynamics of EC gene expression. However, the somites formed in these experimental conditions did not present the same molecular properties as wild-type rostral somites: the Hox signatures of the ectopic somites were similar to somites 8-9 (compatible with a cervical identity) (Dias et al., 2014) and the expression patterns of EC genes did not match those described in the chick occipital somites (Rodrigues et al., 2006). Importantly, somites did not form in a linear array of segments, limiting what can be extrapolated to the physiological setting (Hubaud & Pourquié, 2014).

Our data evidence that *hairy1* gene expression is highly dynamic in primitive streak embryos, prior to the formation of the first somite. This is in agreement with what was previously reported for *hairy2* and *lfng* (Jouve et al., 2002), where five waves of expression were proposed to swap the presumptive PSM before the formation of the first somites. Here, we attempted to determine if EC *hairy1* oscillations take place and assessed their periodicity by performing half-explant cultures with a higher temporal resolution. We could not identify a defined periodicity for *hairy1* oscillations in these early stages, suggesting that either *hairy1* expression is not oscillatory (albeit very dynamic), or that the oscillation period is variable. The time required to form the first five somites is also irregular, which is consistent with EC period variability. As soon as somite formation begins, *hairy1* expression becomes clearly oscillatory, with cycles of expression ranging from 75 to 90 min, closely matching the average time required to form a new morphological somite.

*Her6* expression transitions from fluctuating to oscillatory, mediated by miR-9 negative feedback regulation, as neuronal differentiation occurs (Soto et al., 2020). A similar phenomenon might be taking place in the chick embryo as the first somite is formed. We anticipate that live imaging using a transgenic *hairy1*-reporter chicken model will be key to conclusively elucidate this matter. This was successfully achieved in zebrafish (Riedel-Kruse et al., 2007) and, recently, in mouse embryos (Falk et al., 2022) where the onset of EC oscillations precedes the formation of the very first somite. Here, we combined the classical explant culture approach (Palmeirim et al., 1997) with *in vivo* imaging detection of the formation of two consecutive somite clefts, to assess if periodicity of *hairy1* oscillations and early somite formation are also coupled in the chick model. Using this approach, we were able to definitively establish that the formation of each new somite is accompanied by a complete cycle of *hairy1* expression.

## CONCLUSION

Our work provides novel insights on the spatio-temporal dynamics of early stages of somitogenesis in the chicken embryo. We show that the first somites form sequentially and at a faster rate (75 minutes periodicity), increasing to 90 minutes by 6-somite stage, the time reported for trunk somite formation (Palmeirim et al., 1997). EC *hairy1* gene expression is very dynamic in gastrulation stages, and the time required to recapitulate its expression pattern closely matches the rate of early somite formation. Therefore, here we clearly show that *hairy1* oscillations and somite formation are coupled since the onset of somitogenesis.

## ACKNOWLEDGEMENTS

The authors thank G. Carraco for suggesting the *in vivo* coupling experiment and I. Palmeirim for fruitful discussions and critical reading of the manuscript. The authors thank the support of the Microscopy Unit of ABC-RI, Portuguese Platform for BioImage (reference PPBI-POCI-01-0145-FEDER-022122).

## Notes

**Funding information:** This work was supported by Fundação para a Ciência e Tecnologia, SFRH/BD/146043/2019 to ACMF, PTDC/BEX/BID/5410/2014 to AMJ and RPA, 2021.00238.CEECIND and RISE - LA/P/0053/2020 to ID, by AD-ABC to RPA and TPA and by Município de Loulé.

https://figshare.com/articles/dataset/SomiteExplorerR_Datasets_used_to_infer_individual_somite_period_and_length_using_systematic_measurements_of_the_whole_embryo_segmented_region_/22341070

https://github.com/iduarte/SomiteExplorerR

### Competing Interest Statement

The authors have declared no competing interest.

## REFERENCES

Carraco, G., Martins-Jesus, A. P., & Andrade, R. P. (2022). The vertebrate Embryo Clock : Common players dancing to a different beat. August, 1–21. https://doi.org/10.3389/fcell.2022.944016

Chapman, S. C., Collignon, J., Schoenwolf, G. C., & Lumsden, A. (2001). Improved method for chick whole-embryo culture using a filter paper carrier. Dev Dyn, 220(3), 284–289. https://doi.org/10.1002/1097-0177(20010301)220:3<284::AID-DVDY1102>3.0.CO;2-5

Christ, B., Huang, R., & Wilting, J. (2000). The development of the avian vertebral column. Anatomy and Embryology 2000 202:3, 202(3), 179–194. https://doi.org/10.1007/S004290000114

Christ, Bodo, & Ordahl, C. P. (1995). Early stages of chick somite development. Anatomy and Embryology 1995 191:5, 191(5), 381–396. https://doi.org/10.1007/BF00304424

Dias, A. S., De Almeida, I., Belmonte, J. M., Glazier, J. A., & Stern, C. D. (2014). Somites without a clock. Science, 343(6172), 791–795. https://doi.org/10.1126/science.1247575

Falk, H. J., Tomita, T., Mönke, G., McDole, K., & Aulehla, A. (2022). Imaging the onset of oscillatory signaling dynamics during mouse embryo gastrulation. Development (Cambridge, England), 149(13). https://doi.org/10.1242/dev.200083

Hamburger, V., & Hamilton, H. L. (1992). A series of normal stages in the development of the chick embryo. 1951. Dev Dyn, 195(4), 231–272. http://www.ncbi.nlm.nih.gov/entrez/query.fcgi?cmd=Retrieve&db=PubMed&dopt=Citation&list_uids=1304821

Heinz Herrmann, M. J. B. Schneider, B. J. Neukom, J. A. M. (1951). Quantitative data on the growth process of the somites of the chick embryo: Linear measurements, volume, protein nitrogen, nucleic acids. Journal of Experimental Zoology Part A Ecological Genetics and Physiology, 118(2), 243–268.

Huang, R., Zhi, Q., Patel, K., Wilting, J., & Christ, B. (2000). Contribution of single somites to the skeleton and muscles of the occipital and cervical regions in avian embryos. Anatomy and Embryology 2000 202:5, 202(5), 375–383. https://doi.org/10.1007/S004290000131

Hubaud, A., & Pourquié, O. (2014). Signalling dynamics in vertebrate segmentation. Nature Reviews Molecular Cell Biology, 15(11), 709–721. https://doi.org/10.1038/nrm3891

Jouve, C., Iimura, T., & Pourquie, O. (2002). Onset of the segmentation clock in the chick embryo: evidence for oscillations in the somite precursors in the primitive streak. Development, 129(5), 1107–1117. http://www.ncbi.nlm.nih.gov/pubmed/11874907

Jouve, C., Palmeirim, I., Henrique, D., Beckers, J., Gossler, A., Ish-Horowicz, D., & Pourquie, O. (2000). Notch signalling is required for cyclic expression of the hairy-like gene HES1 in the presomitic mesoderm. Development, 127(7), 1421–1429. http://www.ncbi.nlm.nih.gov/pubmed/10704388

Kimmel, C. B., Ballard, W. W., Kimmel, S. R., Ullmann, B., & Schilling, T. F. (1995). Stages of embryonic development of the zebrafish. Developmental Dynamics, 203(3), 253–310. https://doi.org/10.1002/AJA.1002030302

Palmeirim, I., Henrique, D., Ish-Horowicz, D., & Pourquie, O. (1997). Avian hairy gene expression identifies a molecular clock linked to vertebrate segmentation and somitogenesis. Cell, 91(5), 639–648. https://doi.org/S0092-8674(00)80451-1 [pii]

R Core Team (2017). R: A language and environment for statistical computing. R Foundation for Statistical Computing, Vienna, Austria. https://www.R-project.org/

R Studio Team (2015). RStudio: Integrated Development for R. RStudio, Inc., Boston, MA. http://www.rstudio.com/

Riedel-Kruse, I. H., Müller, C., & Oates, A. C. (2007). Synchrony dynamics during initiation, failure, and rescue of the segmentation clock. Science (New York, N.Y.), 317(5846), 1911–1915. https://doi.org/10.1126/SCIENCE.1142538

Rodrigues, S., Santos, J., & Palmeirim, I. (2006). Molecular characterization of the rostral-most somites in early somitic stages of the chick embryo. Gene Expression Patterns, 6(7), 673–677. https://doi.org/10.1016/j.modgep.2006.01.004

Schröter, C., Herrgen, L., Cardona, A., Brouhard, G. J., Feldman, B., & Oates, A. C. (2008). Dynamics of zebrafish somitogenesis. Developmental Dynamics, 237(3), 545–553. https://doi.org/10.1002/DVDY.21458

Schubert, M., Holland, L. Z., Stokes, M. D., & Holland, N. D. (2001). Three Amphioxus Wnt Genes (AmphiWnt3, AmphiWnt5, and AmphiWnt6) Associated with the Tail Bud: the Evolution of Somitogenesis in Chordates. Developmental Biology, 240(1), 262–273. https://doi.org/10.1006/DBIO.2001.0460

Soto, X., Biga, V., Kursawe, J., Lea, R., Doostdar, P., Thomas, R., & Papalopulu, N. (2020). Dynamic properties of noise and Her6 levels are optimized by miR-9, allowing the decoding of the Her6 oscillator. The EMBO Journal, 39(12), 1–23. https://doi.org/10.15252/embj.2019103558

Tam, P. P. L. (1981). The control of somitogenesis in mouse embryos. Journal of Embryology and Experimental Morphology, 65(Suppl.), 103–128. https://doi.org/10.1242/dev.65.supplement.103

Tenin, G., Wright, D., Ferjentsik, Z., Bone, R., McGrew, M. J., & Maroto, M. (2010). The chick somitogenesis oscillator is arrested before all paraxial mesoderm is segmented into somites. BMC Developmental Biology, 10, 13–20. https://doi.org/10.1186/1471-213X-10-24

