## Supplementary for "SPATIO-TEMPORAL DYNAMICS OF EARLY SOMITE SEGMENTATION IN THE CHICKEN EMBRYO"

### SUPPLEMENTARY MATERIAL

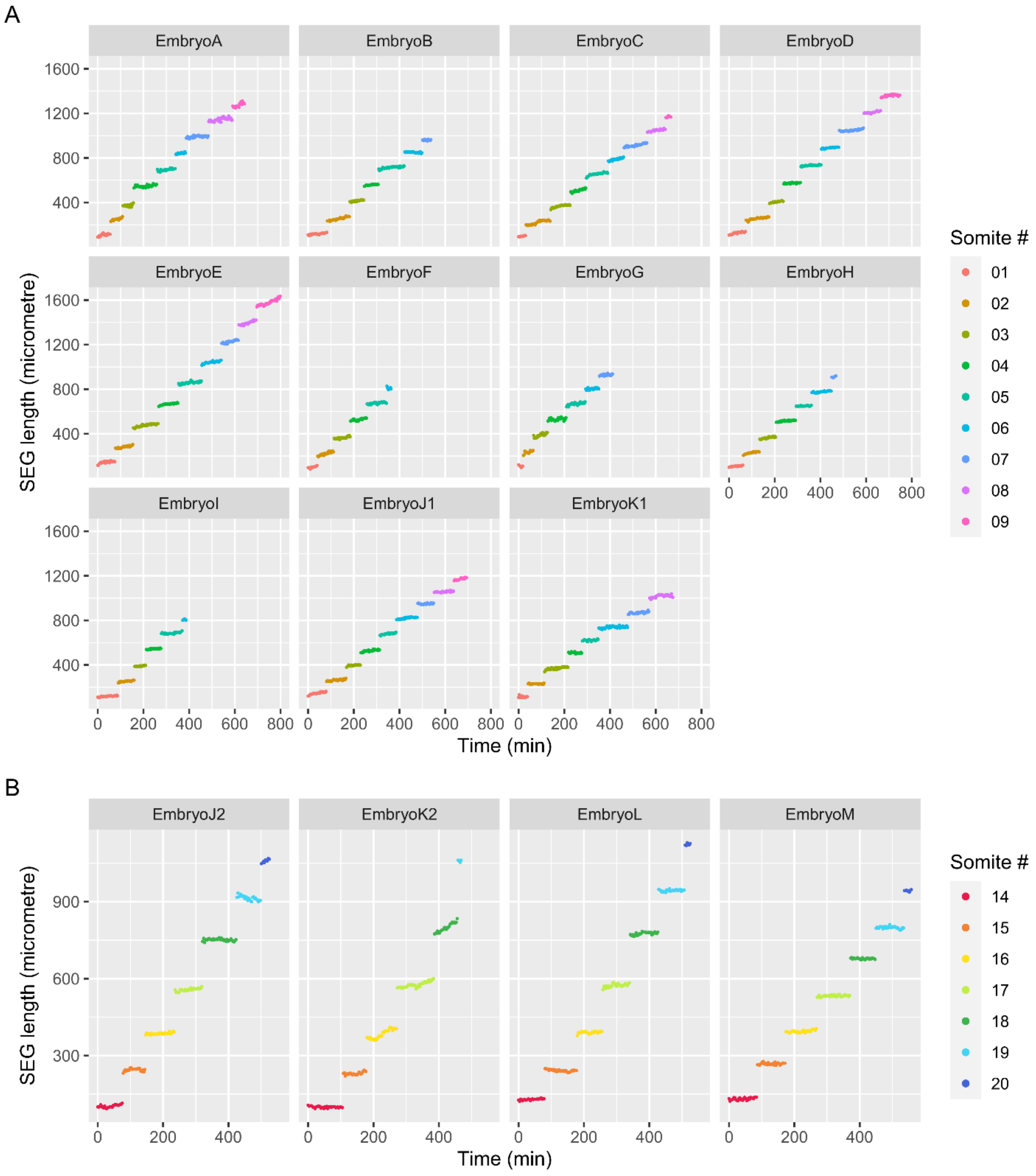


**Supplementary Figure 1 - Length of the segmented region (SEG) of chicken embryos**, from 1 to 9 somites **(A)** and 14 to 20 somites **(B)**, over time. Embryos J and K were cultured continuously from HH7 (1 somite-stage) until HH13+ (20 somite-stage). J1 and K1 correspond to measurements from somites 1-10; J2 and K2 correspond to measurements from somites 14-20.

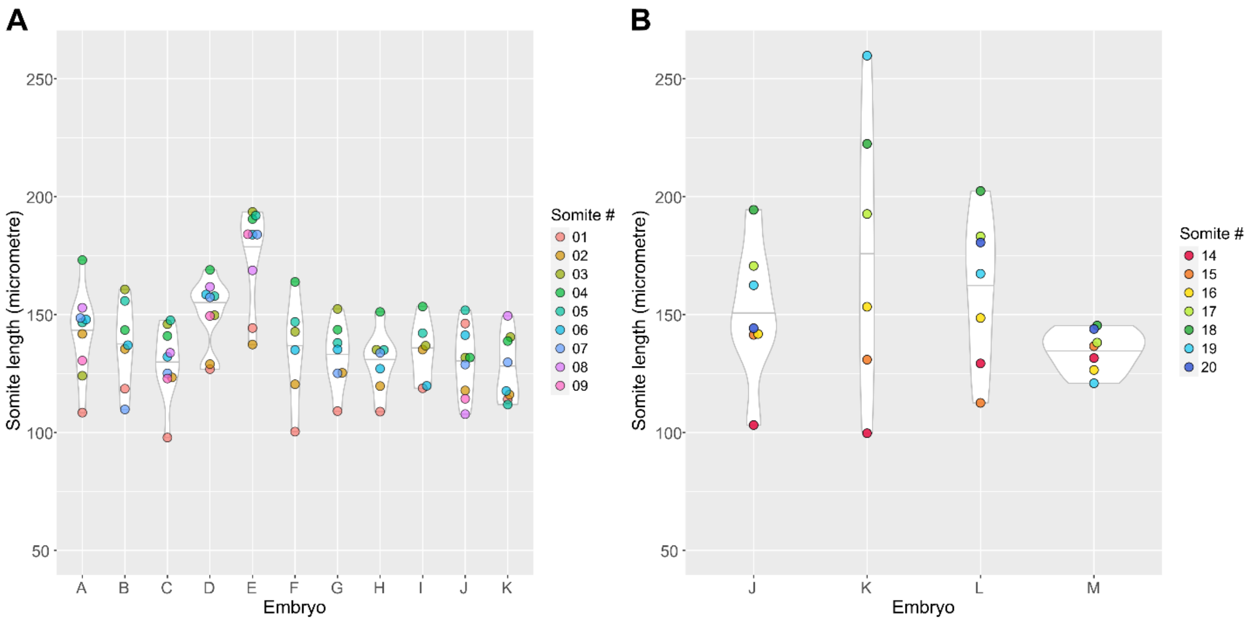


**Supplementary Figure 2 - Somite length per embryo.** Violin-plot distribution of the observed lengths of somites 1-9 **(A)** and 14-20 **(B)** in each of the embryos analysed.

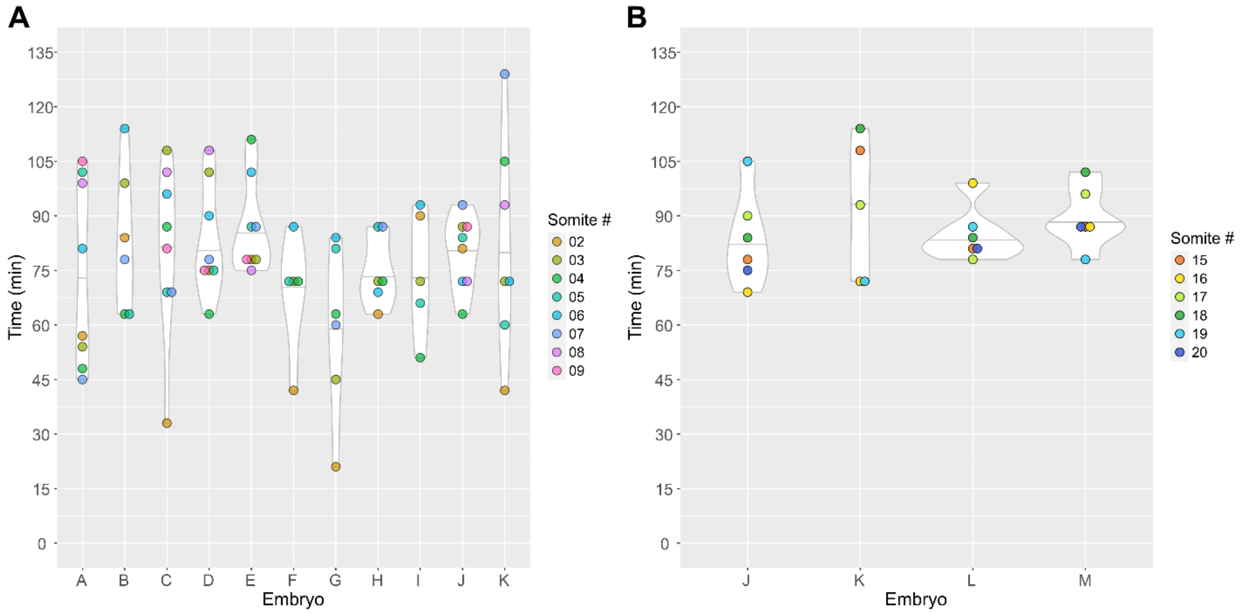


**Supplementary Figure 3 - Somite formation time per embryo.** Violin-plot distribution of the observed periods of somites 2-9 **(A)** and 15-20 **(B)** in each of the embryos analysed.

**Supplementary Table 1 -** Length and Time of formation of chick early somites per embryo analysed. Mean ± standard deviation is shown.

| Embryo | Somites sampled (n) | Length (µm) | Time of Formation (min) |
|---|---|---|---|
| A | 9 | 141.54 ± 18.6 | 73.88 ± 25.7 |
| B | 7 | 137.23 ± 18.43 | 83.5 ± 20.21 |
| C | 9 | 129.95 ± 15.23 | 80.62 ± 23.97 |
| D | 9 | 151.04 ± 14.37 | 83.25 ± 15.36 |
| E | 9 | 175.36 ± 20.97 | 87 ± 13.03 |
| F | 6 | 134.88 ± 22.12 | 69 ± 16.43 |
| G | 7 | 132.67 ± 14.21 | 59 ± 23.52 |
| H | 7 | 130.08 ± 13.38 | 75 ± 9.86 |
| I | 6 | 134.33 ± 13.31 | 74.4 ± 17.42 |
| J1 | 9 | 130.17 ± 14.84 | 79.88 ± 10.01 |
| K1 | 8 | 127.26 ± 14.33 | 81.86 ± 29.28 |
| J2 | 7 | 151.15 ± 28.62 | 83.5 ± 12.79 |
| K2 | 6 | 176.45 ± 59.71 | 91.8 ± 19.63 |
| L | 7 | 160.55 ± 31.98 | 85 ± 7.51 |
| M | 7 | 134.7 ± 8.95 | 89.5 ± 8.36 |

**Embryos J and K were cultured continuously from HH7 (1 somite-stage) until HH13+ (20 somite-stage). J1 and K1 correspond to measurements from somites 1-10; J2 and K2 correspond to measurements from somites 14-20.
